# A hypothalamo-septo-hippocampal circuit for REM sleep-dependent consolidation of social memory

**DOI:** 10.64898/2026.01.13.699230

**Authors:** Tingliang Jian, Wenjun Jin, Mengru Liang, Xiang Liao, Kuan Zhang, Shanshan Liang, Chunqing Zhang, Chao He, Hongbo Jia, Yanjiang Wang, Jian Han, Xiaowei Chen, Han Qin

## Abstract

REM sleep constitutes a critical window for memory consolidation, yet the brain circuits orchestrating this process remain incompletely defined. Here, we identify a lateral supramammillary nucleus (SuM)-medial septum (MS) projection as a REM sleep-specialized pathway essential for hippocampal memory consolidation. Fiber photometry and optrode recordings revealed that lateral SuM-MS projecting neurons were selectively active during REM sleep. REM-specific optogenetic silencing of this projection impaired consolidation of both social and contextual fear memories. Crucially, silencing of its downstream target, the MS-CA2 pathway, during REM sleep selectively disrupted social memory while sparing contextual fear memory. This functional dissection establishes a hypothalamo-septo-hippocampal circuit (lateral SuM-MS-CA2) dedicated to social memory processing, in parallel to the recently-described direct SuM-CA2 pathway. These results also position the SuM as a REM sleep-hub that routes information via parallel septal pathways to consolidate distinct memory modalities.

## Introduction

Sleep is a critical period for memory consolidation, during which newly encoded memories undergo a series of active processes to be strengthened and integrated into long-term memory networks (Diekelmann and Born, 2010, Rasch and Born, 2013, Brodt et al., 2023, Wang et al., 2026). This process does not occur uniformly but is accomplished through the synergistic efforts of both non-rapid eye movement (NREM) and rapid eye movement (REM) sleep (Rial et al., 2023, Falup-Pecurariu et al., 2021). NREM and REM sleep together form a complementary memory processing system essential for transforming transient experiences into lasting, accessible memory (Ackermann and Rasch, 2014, Kaida et al., 2023).

The prevailing concept in the field is that memory consolidation during NREM sleep is critically supported by the reactivation of hippocampal neurons through sharp-wave ripples (SWRs) and neocortical slow-waves (Poe et al., 2010). These coupled oscillatory events are thought to facilitate the redistribution of newly encoded information from the hippocampus to neocortical networks for long-term storage, a process essential for system-level memory consolidation (Born et al., 2006, Buzsaki, 2015, Robinson et al., 2025, Duan et al., 2025). Notably, such SWR-mediated reactivation originates primarily in the CA3 region and has been consistently shown to be necessary for the consolidation of spatial memory (Ego-Stengel and Wilson, 2010, Sasaki et al., 2018). Experimental suppression of SWRs or selective disruption of CA3 neuronal activity during sleep impairs the stabilization of spatial representations, underscoring the functional importance of this mechanism (Girardeau et al., 2009, Ego-Stengel and Wilson, 2010, Roux et al., 2017, Fernandez-Ruiz et al., 2019). In a similar vein, recent work by Oliva et al. has extended this framework by demonstrating that the reactivation of CA2 pyramidal neurons during SWRs is essential for social memory consolidation (Oliva et al., 2016, Oliva et al., 2020, Karaba et al., 2024). These findings not only implicate a previously underappreciated hippocampal subregion in sleep-dependent memory processes but also suggest that distinct cell populations within the hippocampus may support the consolidation of different memory types through a shared SWR-mediated mechanism (Xie et al., 2023).

The REM sleep stage, defined by high-frequency brain activity and vivid dreaming, is considered critical for memory consolidation (Peever and Fuller, 2017). It has been established that the suppression of medial septal (MS) GABAergic neurons during REM sleep leads to a marked decrease in CA1 theta power, resulting in deficits in spatial and contextual memory (Boyce et al., 2016). Complementing this, the work by Kumar et al. revealed a reactivation of adult-born neurons in the dentate gyrus (DG) during REM sleep following contextual fear learning, and this reactivation is essential for memory consolidation (Kumar et al., 2020). Our previous research demonstrated that the hypothalamic supramammillary nucleus (SuM) to hippocampal DG and CA2 pathways are selectively active during REM sleep. We established a causal link through optogenetic interventions, showing that REM-selective silencing of SuM-DG projections impairs spatial and contextual fear memory, whereas similar silencing of SuM-CA2 projections specifically disrupts social memory consolidation (Qin et al., 2022). Furthermore, Liu et al. elucidated how oxytocin modulates the inhibitory balance in the prelimbic cortex to promote social memory consolidation during REM sleep (Liu et al., 2025). Despite these advances, a key question remains: whether additional neural pathways are involved in REM sleep-related social memory consolidation.

SuM and MS are interconnected subcortical hubs that collectively influence a range of behavioral states and cognitive functions. The SuM regulates processes such as episodic memory (Qin et al., 2022, Luo et al., 2025), social and contextual novelty detection (Chen et al., 2020), and wakefulness (Pedersen et al., 2017), whereas the MS, composed of cholinergic, GABAergic, and glutamatergic neurons (Kiss et al., 1990, Hajszan et al., 2004), is critical for hippocampal theta generation (Buzsáki, 2002, Király et al., 2023), memory and the modulation of arousal (Osborne, 1994, An et al., 2021). Building upon this anatomical and functional background, we previously investigated the SuM-MS pathway and found that its activation potently promotes wakefulness (Liang et al., 2023). Furthermore, we identified two functionally distinct subpopulations within this pathway: wake-active and REM-active neurons. Although the wake-active neurons have been causally linked to arousal control, the functional role of the concomitant REM-active neurons remains unknown.

In this study, we employed a comprehensive approach combining optical Ca²⁺ imaging, optrode recordings, and optogenetics to investigate the role of REM sleep in consolidating social memory via a specific hypothalamic-septal-CA2 circuit in mice. We identified a distinct neuronal ensemble within the lateral SuM that projects to the MS and is selectively active during REM sleep. Crucially, REM-selective optogenetic silencing of these SuM-MS projecting neurons impaired both contextual and social memory consolidation. Furthermore, optogenetic inhibition of the downstream MS-CA2 projection during REM sleep selectively disrupted social memory, while sparing contextual fear memory. Together, our findings provide evidence that a population of SuM neurons supports social memory consolidation through its REM sleep-locked activity and targeted influence on the septo-hippocampal route.

## Results

### Lateral SuM-MS projecting neurons are strongly active during REM sleep

SuM neurons have been reported to project to both MS region and hippocampus (Vertes, 1992). We labeled the MS-projecting and hippocampus-projecting SuM neurons by injecting a retrograde AAV vector expressing enhanced green fluorescent protein (EGFP) into MS, and a retrograde AAV vector expressing red fluorescent protein mCherry into hippocampal DG respectively (Figure 1a). We verified the expression area of EGFP in MS and mCherry in hippocampus by post-hoc histology (Figures 1b and 1c). And the EGFP-and mCherry-labeled cell bodies were observed in SuM in serial sections (Figure 1d). The mCherry-labeled neurons were mainly located in the lateral SuM, while the EGFP-labeled ones were distributed in both medial and lateral SuM (Figure 1d). The co-labeling rate of EGFP and mCherry neurons was relatively low, which was less than 5% (Figure 1e). These results suggest that lateral SuM neurons provide projection to both MS and hippocampus.

**Figure 1.**
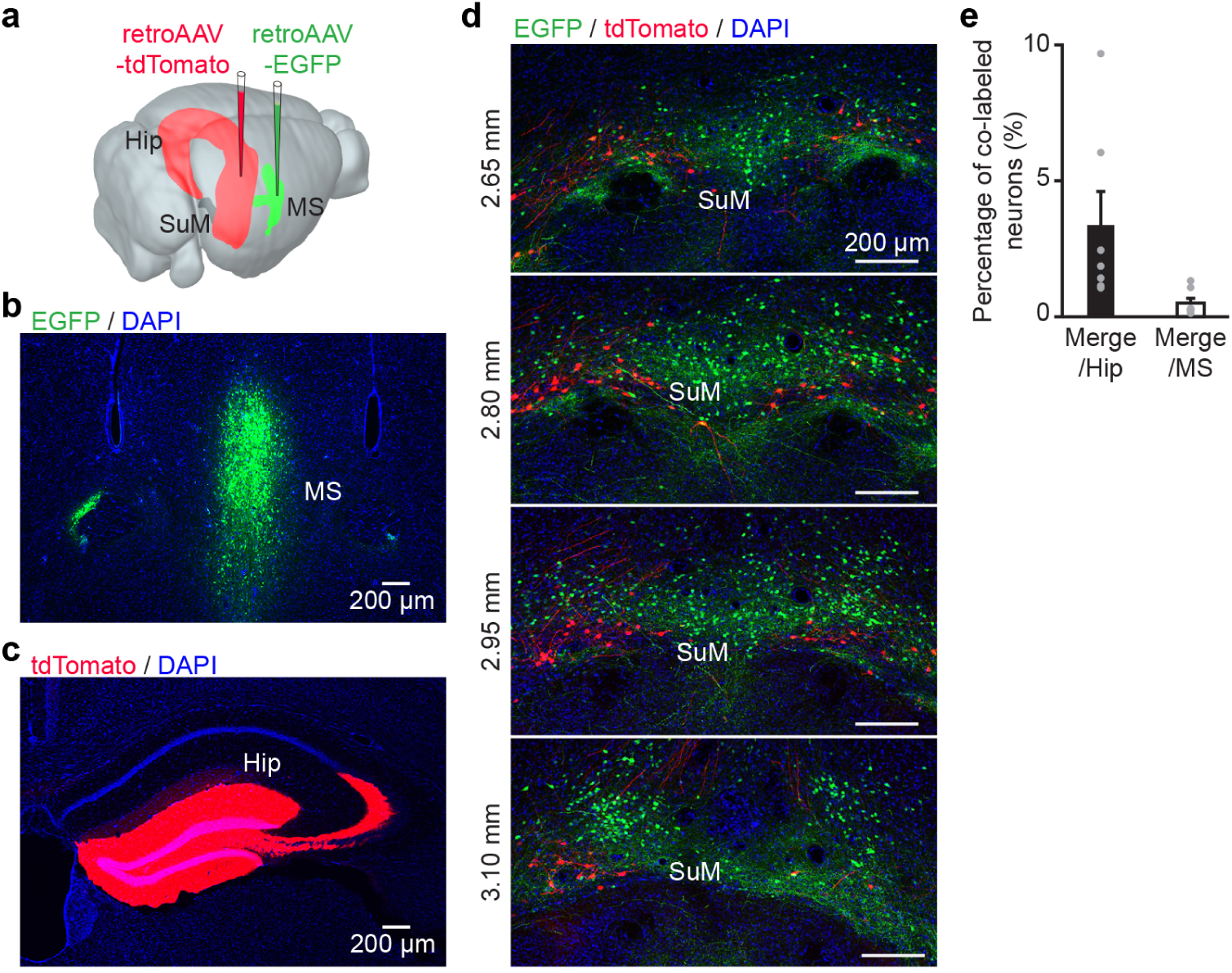
Retrograde labeling of MS-projecting SuM neurons. **a**, Diagram of retroAAV-EGFP injection into MS and retroAAV-tdTomato injection into hippocampus. **b,c,** Post-hoc histological images showing the expression of EGFP at the injection site of MS (b), and the expression of tdTomato at the injection site of hippocampus (c). **d,** Representative serial sections showing the eGFP- and tdTomato-labeled cell bodies in SuM. **e,** Summary of co-labeling proportion of SuM-Hip and SuM-MS projecting neurons, *n* = 6 mice. SuM, supramammillary nucleus; MS, medial septum; Hip, hippocampus.

To determine the neurotransmitter phenotypes of these lateral SuM-MS projecting neurons, we crossed vGlut2-Cre or vGAT-Cre mice with LSL-H2B-GFP reporter mice. A retrograde AAV expressing mRuby was then injected into the MS of these double-transgenic offspring to label the projection neurons (Figure1–figure supplement 1a and 1d). Histological examination revealed that the majority of the mRuby-labeled lateral SuM-MS neurons were co-localized with vGlut2+ (GFP+) nuclei (56.1% ± 2.8%; Figure1–figure supplement 1b and 1c), whereas only a small fraction were positive for vGAT (9.8% ± 2.5%; Figure1–figure supplement 1e and 1f). These anatomical data indicate that the lateral SuM-MS projection is predominantly glutamatergic, consistent with previous studies (Liang et al., 2023, Kesner et al., 2021).

A previous study has demonstrated that Ca^2+^ activity in the axonal terminals of SuM-MS projection was activated during both wakefulness and REM sleep (Liang et al., 2023). However, while these recordings encompassed the entire population of SuM-MS projection (Liang et al., 2023), the activity patterns of those originating specifically from the lateral SuM during the sleep-wake cycle remain uncharacterized. To address this, we employed a circuit-specific fiber photometry system combined with electroencephalogram (EEG) and electromyogram (EMG) recordings to monitor Ca²⁺ dynamics of lateral SuM-MS projecting neurons in freely moving mice. We injected retroAAV-GCaMP6m into MS to selectively express the calcium indicator GCaMP6m in SuM-MS projecting neurons (Figure 2a). Three weeks post-injection, an optical fiber was implanted above the lateral SuM for Ca²⁺ recording, while EEG and EMG electrodes were affixed to the cortical surface and neck muscles, respectively, to monitor behavioral states (Figure 2a). Viral expression and fiber location were confirmed histologically after recordings (Figure 2b). Notably, lateral SuM-MS projecting neurons exhibited higher Ca²⁺ activity during REM sleep compared to NREM sleep and quiet wakefulness (QW, representative trace in Figure 2c; summary in Figure 2d, RMs 1-way ANOVA with Bonferroni post-hoc comparison, F_2,10_ = 19.9, P < 0.001). Furthermore, activity robustly increased during transitions from NREM to REM sleep (Figure 2g) but declined sharply upon transitions from REM sleep to QW (Figure 2h).

**Figure 2.**
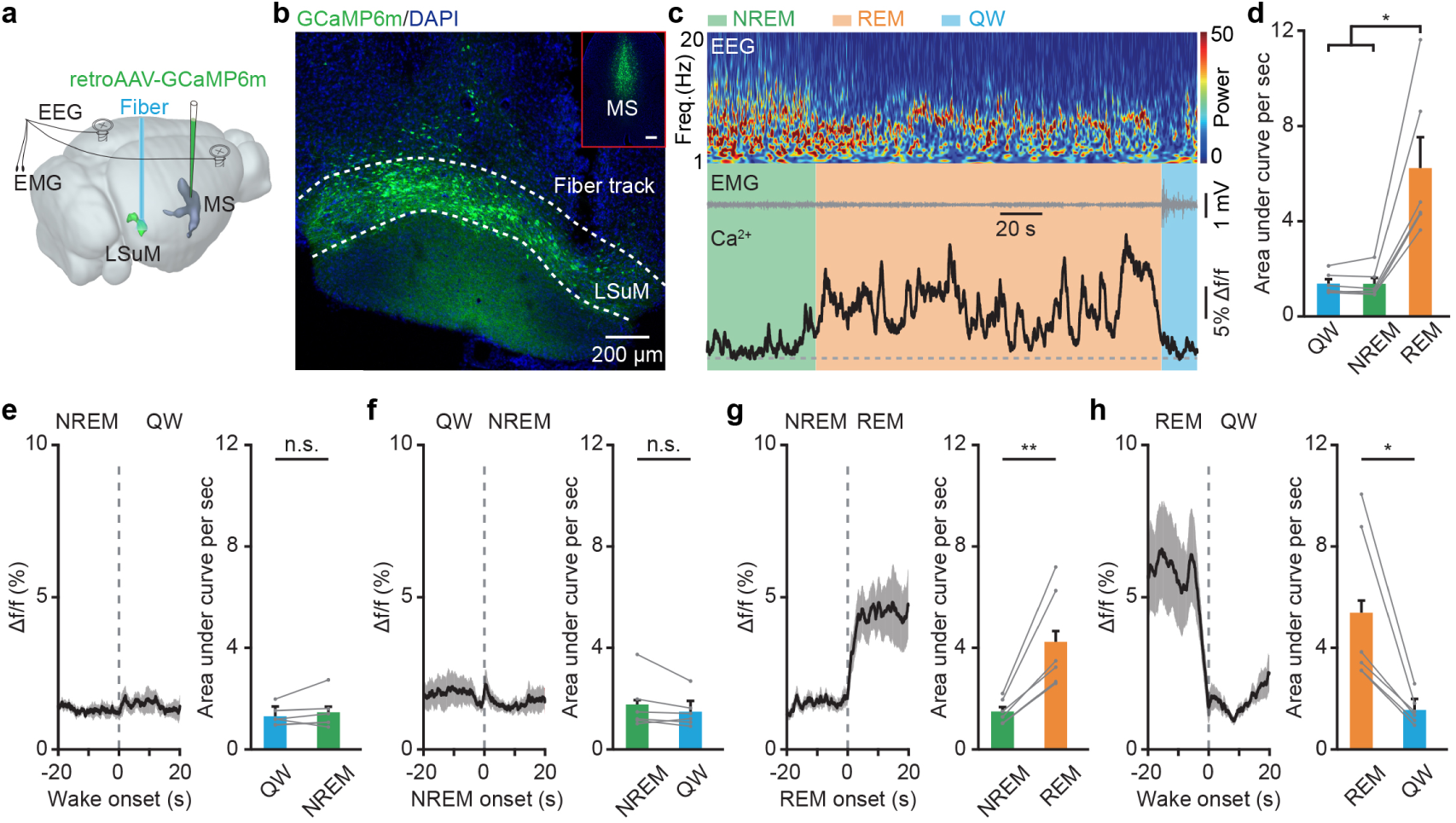
Fiber photometry recording of bulk Ca²⁺ dynamics in lateral SuM-MS projecting neurons across sleep-wakefulness cycle. **a**, Experimental design for virus injection in MS and fiber implantation in lateral SuM. **b,** Post-hoc histological image showing the expression of GCaMP6m at injection site in MS (insert) and in SuM cell bodies, and optical fiber location in lateral SuM. **c,** Ca^2+^ activities in lateral SuM-MS projecting neurons across sleep-wakefulness cycles. Color map indicates power spectrum (μV^2^) of EEG, Freq., frequency. **d,** Summary of the area under the curve per second during wake, NREM sleep, and REM sleep. *n* = 6 mice, RMs 1-way ANOVA with Bonferroni post-hoc comparison. **e–h,** Ca^2+^ activities during brain state transitions: wake-to-NREM (e), NREM-to-wake (f), NREM-to-REM (g), and REM-to-wake (h). *n* = 6 mice, paired *t* test. n.s. *P* > 0.05, \**P* < 0.05, \*\**P* < 0.01. Data are reported as mean ± SEM.

To further characterize the firing patterns of lateral SuM-MS projecting neurons at the single-cell level, we performed optrode recordings across sleep-wake cycles. Specifically, we expressed channelrhodopsin-2 (ChR2) selectively in these neurons by injecting a retrograde Cre-dependent AAV (retroAAV-Cre) into MS and a Cre-dependent AAV encoding DIO-ChR2-mCherry into SuM (Figure 3a). An optrode was then implanted in lateral SuM to identify and record from SuM-MS projecting neurons (Figure 3a; optrode placement shown in Figure 3b). To identify MS-projecting units, we applied blue light pulses (450 nm, 2 Hz, 10 ms duration, 10 mW). Neurons were classified as MS-projecting if they exhibited short-latency, low-jitter spiking with a high success rate and a spike waveform highly correlated with spontaneous spike (mean latency: 3.61 ± 0.27 ms; jitter: 0.73 ± 0.13 ms; success rate: 90% ± 2%; correlation coefficient: 0.91 ± 0.02; n = 29 neurons; example neurons in Figure 3c and 3d; summary in Figures 3e–3g).

**Figure 3.**
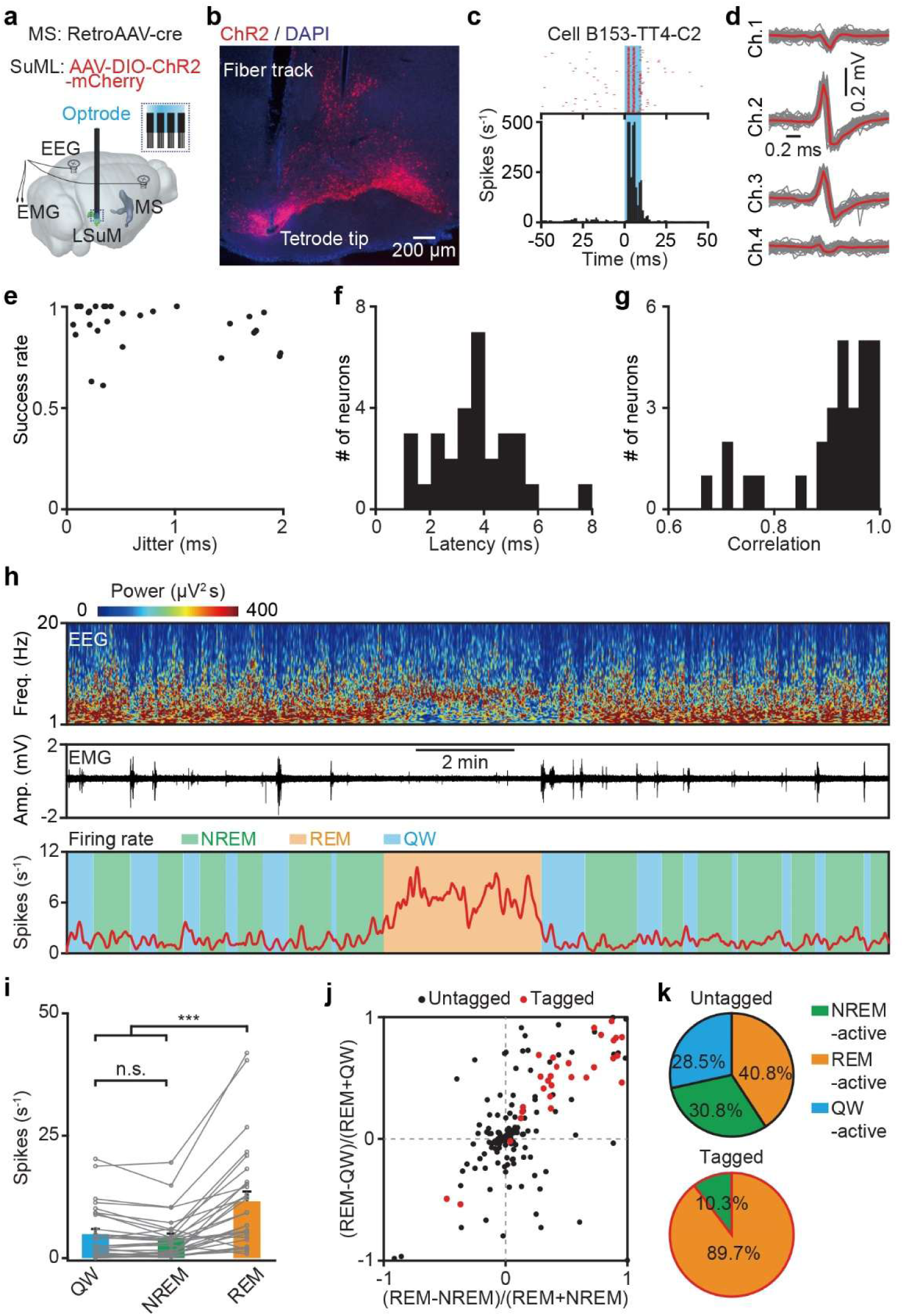
Optrode recording of single-unit activity in lateral SuM-MS projecting neurons across sleep-wakefulness cycle. **a**, Experimental design for virus injection and optrode implantation in the lateral SuM-MS projection. **b,** Representative image showing virus expression and locations of fiber and tetrode tips in lateral SuM. **c,** Histogram of stimulus time for spikes in a representative lateral SuM-MS projecting neuron (Cell#B153-TT4-C2). **d,** Waveforms of spontaneous (grey) and light-induced (red) spikes from the unit in (c). **e,** Success rate versus temporal jitter of the first light-induced spikes for all recorded lateral SuM-MS projecting neurons, *n* = 29 neurons from 15 mice. **f,g,** Distributions of latencies of the first light-induced spikes (f), and correlation coefficients between light-induced spikes and spontaneous spikes (g) for all recorded lateral SuM-MS projecting neurons, *n* = 29 neurons from 15 mice. **h,** Firing rate of the neuron in (c) across sleep-wakefulness cycles. Freq., frequency. QW, quiet wakefulness. **i,** Firing rates of 29 tagged lateral SuM-MS projecting neurons during three different states. Friedman test with Bonferroni post-hoc comparison. **j,** Firing rate modulation of lateral SuM-MS projecting neurons (red, *n* = 29 neurons from 15 mice) and non-MS-projecting lateral SuM neurons (black, *n* = 130 neurons from 15 mice). **k,** Distribution of REM-active, NREM-active and QW-active neurons of MS-projecting and non-MS-projecting lateral SuM neurons. n.s. *P* > 0.05, \*\*\**P* < 0.001. Data are reported as mean ± SEM.

We found that the majority of lateral SuM-MS projecting neurons (26 of 29 units from 15 mice) exhibited a significant increase in firing rates upon transition from NREM to REM sleep, followed by a marked decrease when transitioning from REM sleep to QW (example in Figure 3h; summary in Figure 3i; REM: 11.6 ± 2.0 Hz, NREM: 4.2 ± 0.9 Hz, QW: 4.9 ± 1.0 Hz; Friedman test with Bonferroni correction, χ^2^ = 29.8, p < 0.001). Analysis of firing patterns across tagged and untagged neurons recorded with the same tetrodes revealed that approximately 90% of tagged neurons were REM-active, with the remaining 10% active primarily during NREM sleep. In contrast, among untagged neurons (n = 130), about 41% showed high activity during REM sleep, while the rest were predominantly active during NREM sleep (31%) or QW (28%; Figures 3j and 3k). Collectively, these results indicate that lateral SuM-MS projecting neurons are highly active during REM sleep, but not during QW.

### Silencing of Lateral SuM-MS projecting neurons during REM sleep impairs both social and contextual memory

Previous studies have revealed that strong activation of SuM-CA2 projection during REM sleep promotes social memory consolidation (Qin et al., 2022). It remains unknown whether the similarly strong activation of lateral SuM-MS projection contributes to hippocampal memory consolidation during REM sleep. To address this, we tested hippocampus-dependent memory after REM sleep-specific silencing of the lateral SuM-MS projection using projection-targeted optogenetics. Specifically, the inhibitory opsin Guillardia theta anion-conducting channelrhodopsin 1 (GtACR1, (Govorunova et al., 2015) was expressed in lateral SuM-MS projecting neurons via retrograde AAV-Cre injected into the MS and Cre-dependent AAV-DIO-GtACR1-mCherry delivered into the lateral SuM (Figures 4a and 4b). We first investigated whether silencing these neurons affects hippocampal CA1 activity during REM sleep by recording local field potentials (LFP). Photoinhibition did not alter LFP power in the dorsal CA1 across the 0–30 Hz frequency range, which includes delta, theta, alpha, and beta rhythms (Figures 4c, d). This contrasts with earlier reports in which silencing other neuronal populations, MS GABAergic neurons (Boyce et al., 2016) or SuM-DG projecting neurons (Qin et al., 2022), significantly reduced theta rhythm in the dorsal CA1.

**Figure 4.**
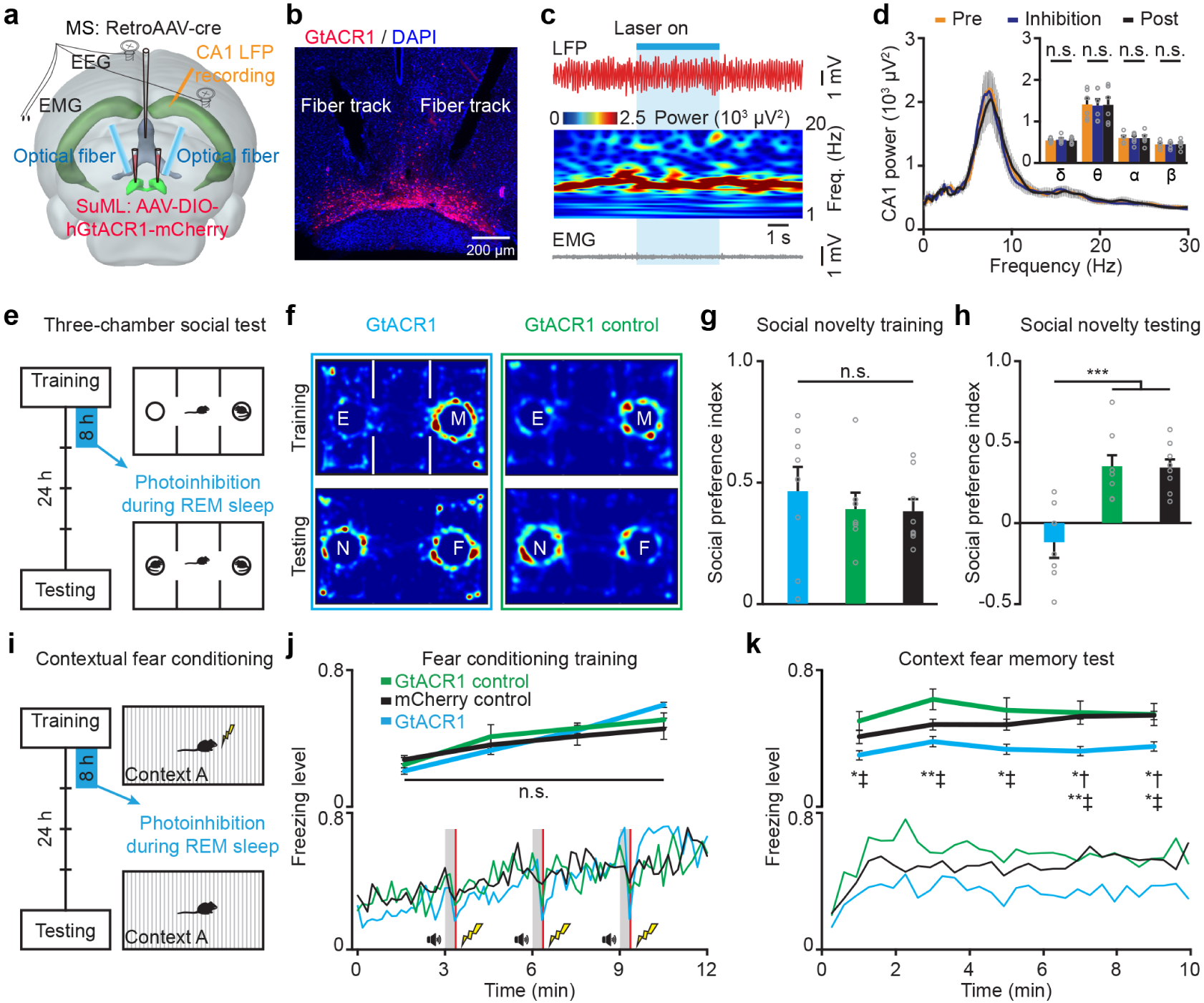
Optogenetic silencing of lateral SuM-MS projecting neurons during REM sleep impairs both social and contextual fear memory. **a–d**, Effects of silencing lateral SuM-MS projecting neurons on CA1 local field potential (LFP) activity during REM sleep. **a,** Experimental design for retrograde labeling of lateral SuM-MS projecting neurons by GtACR1-mCherry, fiber implantation above SuM, and LFP recording in CA1. **b,** Representative image showing virus expression and fiber tracks in lateral SuM. **c,** Representative EEG showing the effects of silencing lateral SuM-MS projecting neurons on CA1 LFP during REM sleep. Freq., frequency. **d,** CA1 LFP power spectral analysis pre-, during, and post-photoinhibition. RMs 1-way ANOVA test, *n* = 6. **e,** Protocol for three-chamber social memory task. E, empty; M, mouse; N, novel; F, familiar. **f,** Representative heatmaps of distribution of time in three-chamber task. **g,h,** Summary of social preference indexes in training (g) and testing (h) of the social novelty task. 1-way ANOVA with Bonferroni post hoc comparison, GtACR1 group *n* = 8, GtACR1 control *n* = 7, mCherry control *n* = 8. **i,** Protocol for optogenetic silencing and contextual fear memory. **j,k,** Freezing levels during training (j) and testing (k) phases. RMs 2-way ANOVA with Bonferroni post-hoc comparison, GtACR1 group *n* = 7, GtACR1 control *n* = 8, mCherry control *n* = 8; †, GtACR1 v.s. mCherry control; ‡ GtACR1 v.s. GtACR1 control. n.s. *P* > 0.05, \**P* < 0.05, \*\**P* < 0.01, \*\*\**P* < 0.001. Data are reported as mean ± SEM.

We next examined the effect of REM sleep-selective silencing of lateral SuM-MS projection on social memory consolidation using the three-chamber social memory test (Hitti and Siegelbaum, 2014). During the training phase, subject mice were allowed to freely explore two chambers for 20 min: one containing an unfamiliar stimulus mouse and the other empty (Figure 4e). All groups showed a significant preference for the chamber with the stimulus mouse during training (Figure 4f top; 201.1 ± 20.9 s vs. 76.7 ± 7.8 s; P < 0.001, Wilcoxon signed-rank test, n = 23 mice). A social preference index was calculated based on cumulative exploration time in the two chambers. No significant difference in this index was observed among the GtACR1, mCherry control, and GtACR1 control groups during training (Figures 4g; P = 0.71, 1-way ANOVA test). During the first 8 hours post-training (the memory consolidation phase), optogenetic silencing was delivered via implanted optical fibers to lateral SuM-MS projecting neurons. In both the GtACR1 group and the mCherry control group, blue light illumination began approximately 6 seconds after REM sleep detection and continued until the end of each REM episode. In the GtACR1 control group (which also expressed GtACR1 in lateral SuM-MS neurons), light was delivered during NREM sleep or QW for durations matched to the total REM-light time in the experimental group.

On day 2 of testing, mice freely explored a chamber containing a familiar mouse and another chamber with a novel mouse (Figure 4e). As conspecifics typically prefer novel social interactions, control groups were expected to show a novelty preference. The GtACR1 group, however, exhibited no significant preference between the novel and familiar mice (Figure 4f bottom left; 150.1 ± 19.5 s vs. 193.4 ± 24.5 s; P = 0.2, Wilcoxon signed-rank test, n = 8 mice). In contrast, both the GtACR1 control group (Figure 4f bottom right; 178.8 ± 51.3 s vs. 85.4 ± 24.3 s; P = 0.02, n = 7 mice) and the mCherry control group (265.5 ± 31.3 s vs. 124.1 ± 10.6 s; P = 0.01, n = 8 mice) retained a clear preference for the novel mouse. Consistent with these findings, the social preference index was significantly lower in the GtACR1 group compared to both control groups (Figure 4h; GtACR1 vs. GtACR1 control, P < 0.001; vs. mCherry control, P < 0.001; 1-way ANOVA with Bonferroni post hoc comparison, F_2,22_ = 13.5, P < 0.001).

We further employed contextual fear conditioning to investigate how silencing lateral SuM-MS projecting neurons during REM sleep affects contextual memory consolidation. First, mice were trained via contextual fear learning on day 1 (Figures 4i). During training, mice showed a significant high freezing level after three paired training (Figure 4j; trained 0.55 ± 0.02 vs. baseline 0.24 ± 0.02; P < 0.001, paired *t*-test, n = 23 mice), with no significant difference in freezing level among the GtACR1, mCherry control and GtACR1 control groups (Figure 4j; P = 0.4, 1-way ANOVA). Within the first 8 hours after training, optogenetic inhibition was delivered to lateral SuM-MS projecting neurons selectively during REM sleep (GtACR1 and mCherry control groups) or during NREM/QW (GtACR1 control group). At 24 hours post-training, mice were re-exposed to the conditioned context for 10 min. The GtACR1 group exhibited significantly lower freezing levels compared to both the mCherry control group and the GtACR1 control group (Figure 4k; RMs 2-way ANOVA with Bonferroni post hoc comparison, F_2,20_ = 8.0, P = 0.003). Together, these results demonstrate that the lateral SuM-MS projection plays a critical role in consolidating both contextual and social memory during REM sleep.

### Silencing of MS-CA2 projecting neurons during REM sleep impairs social but not contextual memory

Previous studies have established the involvement of MS GABAergic neurons in REM sleep-dependent contextual memory consolidation (Boyce et al., 2016). In addition, inhibition of the projection from MS to CA2 has been shown to disrupt social memory encoding (Wu et al., 2021). However, whether the MS-CA2 projection contributes to social memory consolidation remains unclear. To investigate this, we specifically labeled MS-CA2 projecting neurons with GtACR1 by injecting retroAAV-Cre into CA2 and AAV-DIO-GtACR1-mCherry into MS (Figure 5a). Post-hoc histology confirmed the retroAAV-Cre injection site in CA2 (Figure 5b), as well as the viral expression and optical fiber placement in MS (Figure 5c). We first performed the three-chamber social test to assess the effect of REM-selective silencing of MS-CA2 projection on social memory. Overall, mice spent significantly more time interacting with the social stimulus chamber than the empty chamber (Figure 5d top for example and Figure 5e for summary) in the training phase in all groups. We then applied optogenetic silencing specifically during REM sleep in the 8-hour period following training. When retested 24 hours later with a familiar versus a novel mouse, the GtACR1 group exhibited a significantly reduced social preference index compared to both control groups (Figure 5f; GtACR1: 0.02 ± 0.02, GtACR1 control: 0.26 ± 0.04, mCherry control: 0.28 ± 0.04; 1-way ANOVA with Bonferroni post hoc comparison, F_2,28_ = 18.7, P < 0.001; GtACR1 vs. GtACR1 control: P < 0.001, GtACR1 vs. mCherry control: P < 0.001).

**Figure 5.**
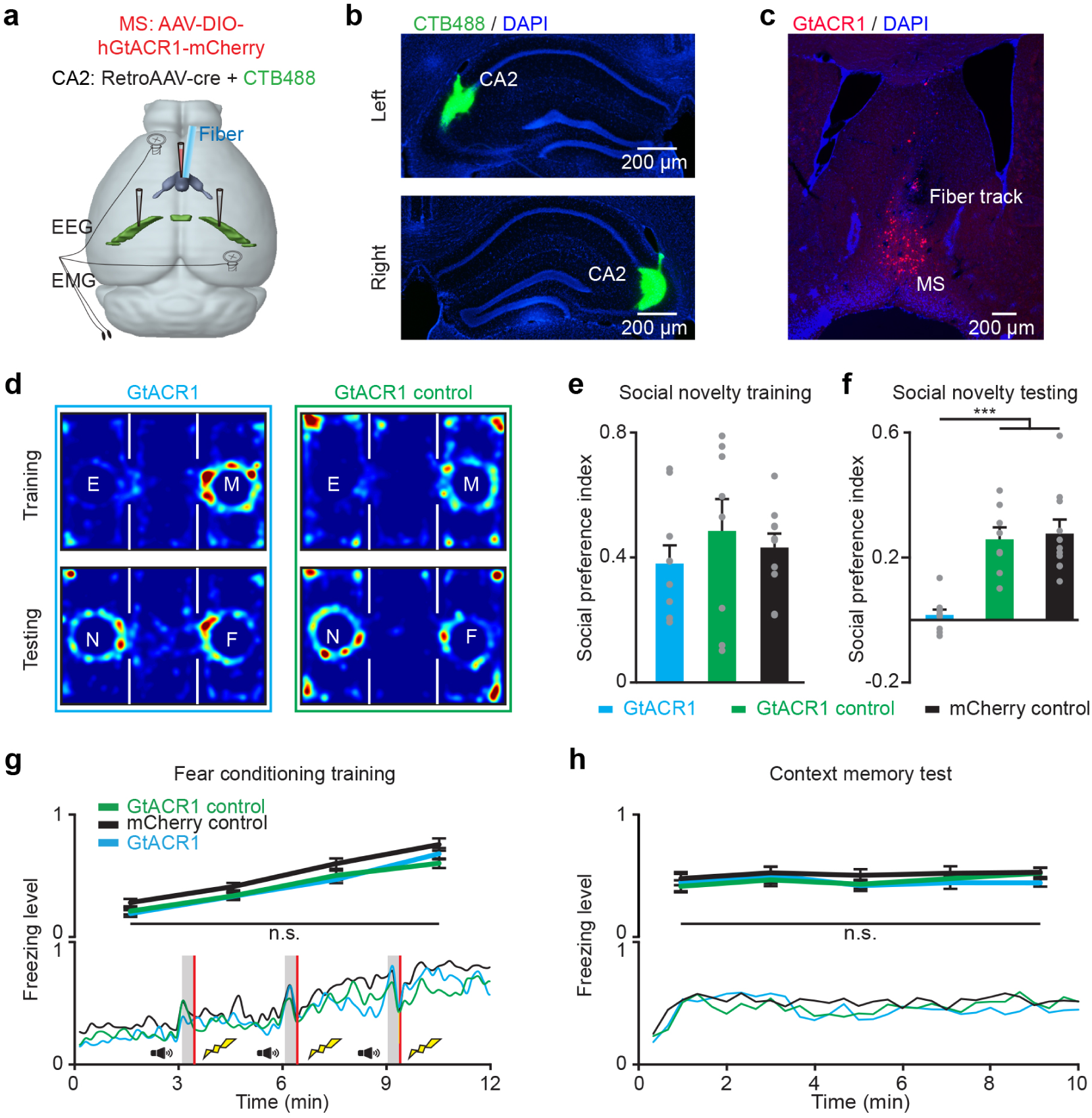
Optogenetic silencing of MS-CA2 projecting neurons during REM sleep impairs social but not contextual fear memory. **a**, Experimental design for retrograde labeling of MS-CA2 projecting neurons by GtACR1-mCherry, and fiber implantation above MS. **b,** Representative images showing the injection site of retroAAV-Cre mixed with CTB488 in CA2. **c,** Representative image showing viral expression and the fiber track in MS. **d,** Representative heatmaps of distribution of time in three-chamber task. **e,f,** Summary of social preference indexes in training (e) and testing (f) of the social novelty task. 1-way ANOVA with Bonferroni post hoc comparison; GtACR1 group *n* = 10, GtACR1 control *n* = 8, mCherry control *n* = 11. **g,h,** Freezing levels during training (g) and testing (h) phases. RMs 2-way ANOVA test; GtACR1 group *n* = 8, GtACR1 control group *n* = 10, mCherry control *n* = 11. n.s. *P* > 0.05, \*\*\**P* < 0.001. Data are reported as mean ± SEM.

We next examined the effect of silencing MS-CA2 projections during REM sleep on contextual fear memory. Mice underwent contextual fear conditioning (Figure 5g), after which MS-CA2 neuronal activity was optogenetically silenced during REM sleep as previously described. During memory retrieval on day 2, no significant differences in freezing level were observed among the three groups (Figure 5h, RMs 2-way ANOVA, P = 0.5). Together, these results demonstrate that the MS-CA2 projection, as a downstream pathway of the SuM-MS circuit, is essential for REM sleep-dependent consolidation of social memory but not contextual memory, thereby completing the picture of a dedicated hypothalamo-septal-hippocampal circuit for sleep-dependent social memory processing.

### Anatomical topology of SuM-recipient MS neurons projecting to CA2

To explore the potential local circuit organization linking the lateral SuM-MS and MS-CA2 pathways, we first sought to identify the neurotransmitter phenotypes of MS neurons receiving direct SuM inputs. To label these cells, we injected AAV2/1-FlpO into lateral SuM and AAV-fDIO-EGFP into MS to anterogradely label SuM-recipient MS neurons. In separate cohorts, DIO-mCherry was injected into MS of vGAT-Cre (Figure5–figure supplement 1a) or vGlut2-Cre (Figure5–figure supplement 1d) mice to label GABAergic or glutamatergic neurons, respectively, while cholinergic neurons were identified via CHAT immunohistochemistry (Figure5–figure supplement 1g). Using these approaches, we found that 54.8% ± 5.3% of these SuM-recipient MS neurons were GABAergic (vGAT^+^ Figure5–figure supplement 1b and 1c), whereas 16.6% ± 1.1% were glutamatergic (vGlut2^+^ Figure5–figure supplement 1e and 1f), with a minor fraction (2.4% ± 0.7%) being cholinergic (CHAT^+^ Figure5–figure supplement 1h and 1i).

We next examined whether these SuM-recipient MS neurons send direct projections to CA2. Using an anterograde tracing strategy (injecting AAV2/1-Cre into lateral SuM and AAV-DIO-EGFP into MS), we observed EGFP-labeled axonal terminals within the CA2 region (Figure5–figure supplement 2). Together, these anatomical data indicate that the SuM-recipient MS neurons are highly heterogeneous, comprising both GABAergic and glutamatergic populations, and that a subset of SuM-recipient MS neurons directly projects to CA2.

## Discussion

Previous work has highlighted the role of SuM-DG and SuM-CA2 projections in REM sleep-dependent memory consolidation (Qin et al., 2022). In contrast, the functional role of SuM-MS pathway is less clear. While it has been implicated in promoting wakefulness (Liang et al., 2023, Duan et al., 2025, Li et al., 2025b, Li et al., 2025a), the contribution of its REM-active neuronal subpopulation remains undefined. Here, by combining recording and manipulation of the lateral SuM-MS subpopulation, we show that it is specifically activated during REM sleep and, together with the downstream MS-CA2 projection, serves as a critical circuit for social memory consolidation. These findings suggest that the SuM may orchestrate distinct aspects of sleep-related cognition through functionally specialized subcircuits.

Our results further delineate a functional dissociation within the supramammillary-septo-hippocampal system. While the MS-DG projection is known to consolidate contextual fear memory by gating dentate gyrus granule cell activity (Boyce et al., 2016, Delorme et al., 2021), and the MS-CA2 projection is essential for social memory encoding (Wu et al., 2021), whether the latter also contributes to social memory consolidation remained unclear. Here, by REM sleep-selective inhibition of the MS-CA2 pathway, we demonstrate its necessary role in consolidating social memory during REM sleep. Collectively, these findings suggest that newly encoded social information in CA2 and contextual information in DG may be reactivated and consolidated both through parallel circuits: the direct supramammillary-hippocampal route (Qin et al., 2022) and the indirect supramammillary-septo-hippocampal pathway (Kiss et al., 2000).

Notably, we found that the SuM-MS pathway did not modulate hippocampal theta rhythms in dorsal CA1, in contrast to the established roles of the SuM-DG projection (Qin et al., 2022) and MS GABAergic neurons in theta generation (Boyce et al., 2016). How can this be reconciled with our anatomical finding that a large proportion of SuM-recipient MS neurons are GABAergic? One likely explanation is the profound heterogeneity of the MS GABAergic population. While global manipulation of MS GABAergic neurons disrupts theta generation (Boyce et al., 2016), the specific subpopulation receiving SuM inputs may represent local interneurons that regulate intra-septal microcircuits, rather than directly serving as hippocampal theta pacemakers. How might this circuit support memory consolidation if not through global theta synchrony? First, given that MS-CA2 projecting neurons are predominantly cholinergic (Wu et al., 2021), the SuM-MS circuit may regulate local acetylcholine release in CA2, thereby altering the neurochemical milieu required for synaptic plasticity. Second, it may drive synapse-specific plasticity at MS-CA2 glutamatergic synapses (Wu et al., 2021), directly strengthening the specific inputs that encode social information. Collectively, these observations suggest that the SuM-MS circuit supports memory consolidation through highly localized, theta-independent mechanisms—such as tuning the local neuromodulatory tone or synapse-specific plasticity within specific hippocampal subfields during REM sleep (Malenka and Bear, 2004, Aime et al., 2022).

Methodologically, the pathway-specific tools employed here were crucial for delineating functional heterogeneity within the SuM-MS pathway. From an anatomical perspective, the SuM can be subdivided into medial and lateral subregions. These subregions differ in their cellular composition and efferent-afferent circuit organization, which likely underlies their distinct physiological roles (Vertes, 1992, Pan and McNaughton, 2004). Our findings reveal a notable regional specialization: lateral SuM-MS projectors are predominantly REM-active (∼90%) and lack detectable wake-active populations, in contrast to the mixed wake/REM profile previously described for medial projectors (∼43% REM-active, ∼57% wake-active, (Liang et al., 2023). Functionally, this segregation suggests that the medial SuM-MS pathway may integrate general arousal or affective signals to promote wakefulness, whereas the lateral SuM-MS pathway might specialize in routing REM-specific mnemonic or emotional information during sleep. This suggests that REM-active neurons, particularly in the lateral SuM, are key drivers of the observed REM-related memory effects, although contributions from medial SuM REM-active or wake-active subgroups cannot be excluded. Consequently, future work should refine methods to selectively manipulate REM-active SuM-MS neurons and systematically compare the roles of medial versus lateral SuM subregions in diverse behaviors, including arousal, locomotion, and memory.

How the SuM-MS and MS-CA2 pathways functionally interact to consolidate social memory remains an open question. Our anatomical tracing reveals a potential local microcircuit linking these structures: while the SuM-MS projection is predominantly glutamatergic, the SuM-recipient MS neurons are highly heterogeneous, comprising both GABAergic and glutamatergic populations (Figure5–figure supplement 1). Given that MS neurons projecting to CA2 are primarily cholinergic and glutamatergic (Wu et al., 2021), and that glutamatergic synapses at the MS-CA2 projection induce plasticity essential for social memory, two distinct functional models can be proposed. First, a direct serial pathway may exist where a small subset of glutamatergic SuM-recipient MS neurons projects to CA2 (Figure5–figure supplement 2), potentially driving REM sleep-dependent social memory consolidation. Second, the GABAergic SuM-recipient MS neurons may act as local interneurons, modulating the activity of CA2-projecting cholinergic or glutamatergic MS neurons via intra-septal microcircuits, thereby indirectly influencing social memory consolidation in CA2. Furthermore, since our current manipulations targeted projection-defined populations, future studies employing intersectional genetic strategies will be necessary to target molecularly defined MS subpopulations and precisely test these microcircuit models. Additionally, expanded behavioral paradigms (e.g., social hierarchy) will help determine whether this circuit broadly regulates social cognition or is specific to novelty-based social memory.

In summary, we reveal a supramammillary-septo-hippocampal circuit that is selectively active during REM sleep and necessary for social memory consolidation. Building on prior work, our findings support a model in which the SuM functions as a REM-hub, directing information via specialized downstream pathways to consolidate distinct memory types. This provides a novel framework for understanding how subcortical-hippocampal interactions during REM sleep govern memory consolidation.

**Figure 1–figure supplement 1.**
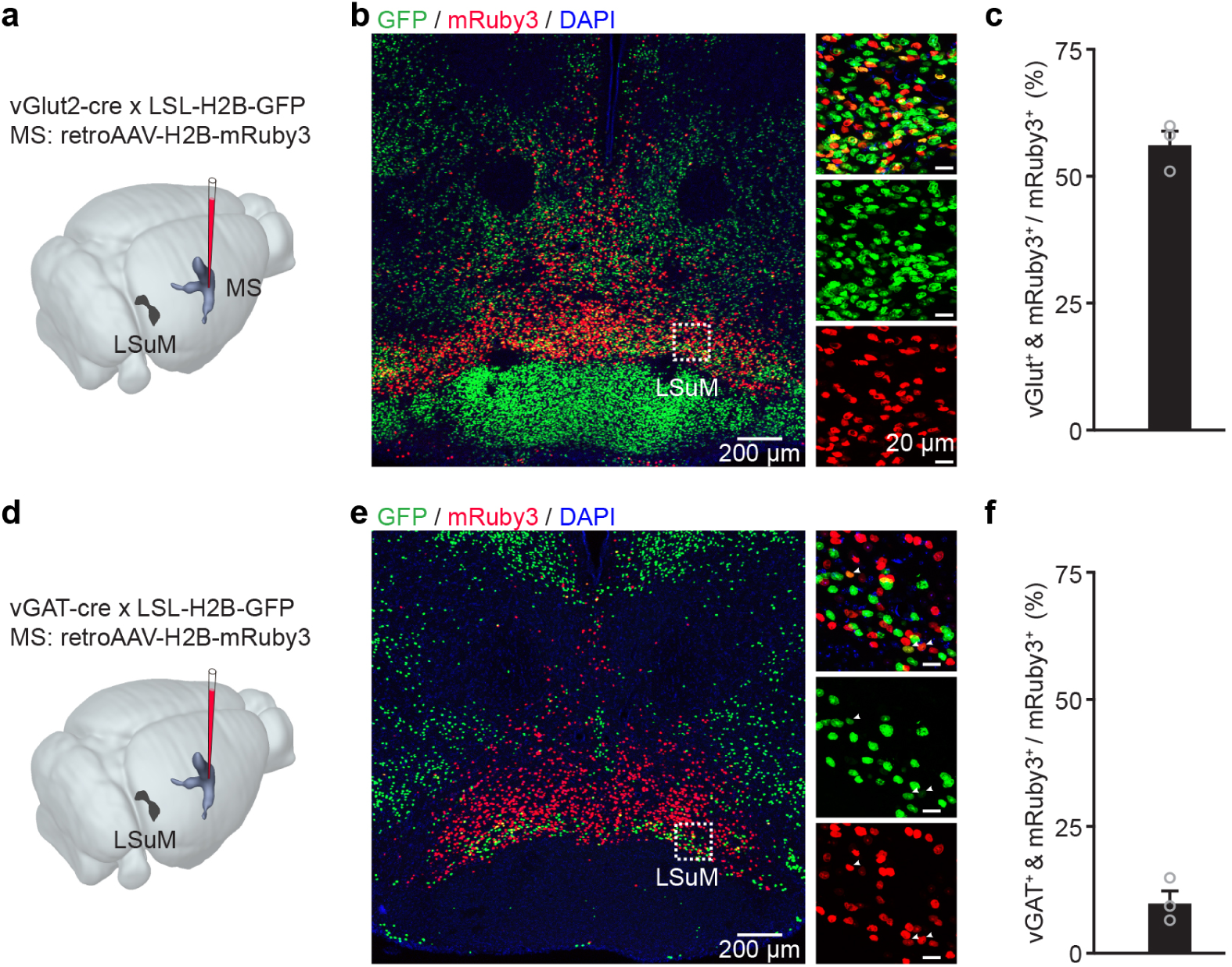
Anatomical validation of the neurotransmitter phenotypes of SuM-MS projecting neurons. **a**, Experimental design for retrograde labeling of lateral SuM-MS projecting neurons by mRubys, and labeling of vGlut2^+^ neurons by crossing of vGlut2-Cre mice with LSL-H2B-GFP reporter mice. **b**, Representative confocal images showing GFP (vGlut2^+^) and mRuby (retrograde) double-labeled cell bodies in the SuM. **c,** Summary of co-labeling proportion of SuM-MS projecting and vGlut2^+^ neurons, *n* = 3 mice. **d**, Experimental design for retrograde labeling of lateral SuM-MS projecting neurons by mRubys, and labeling of vGAT^+^ neurons by crossing of vGAT-Cre mice with LSL-H2B-GFP reporter mice. **e**, Representative confocal images showing GFP (vGAT^+^) and mRuby (retrograde) double-labeled cell bodies in the SuM. **f,** Summary of co-labeling proportion of SuM-MS projecting and vGAT^+^ neurons, *n* = 3 mice. LSuM, lateral supramammillary nucleus; MS, medial septum.

**Figure5–figure supplement 1.**
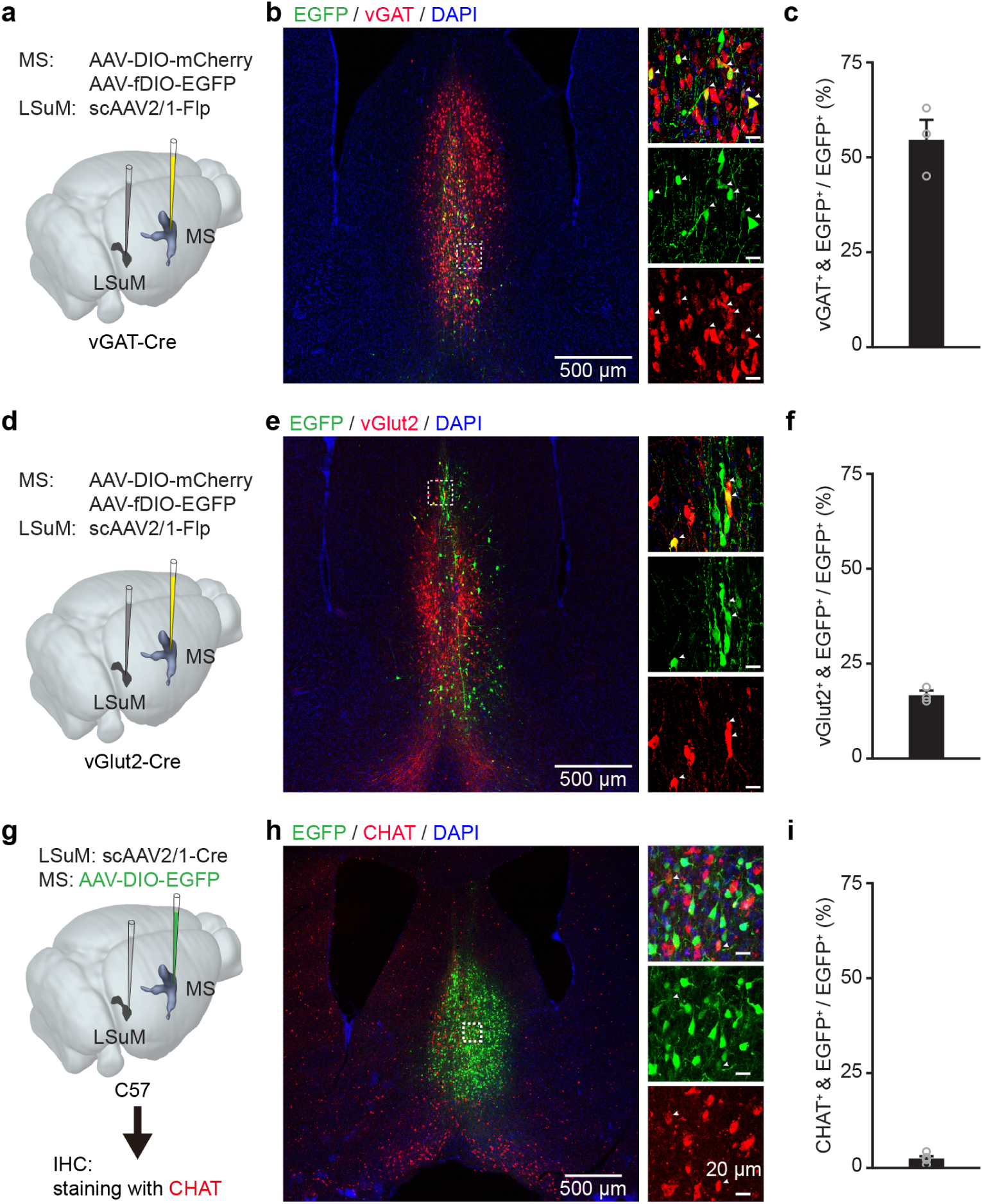
Anatomical identification of the neurotransmitter phenotypes of SuM-recipient MS neurons. **a**, Experimental design for anterograde labeling of SuM-recipient MS neurons by EGFP, and labeling of vGAT^+^ neurons by injection of DIO-mCherry in vGAT-Cre mice. **b**, Representative confocal images showing mCherry (vGAT^+^) and EGFP (anterograde) double-labeled cell bodies in MS. **c**, Summary of co-labeling proportion of SuM-recipient and vGAT+ neurons in MS, *n* = 3 mice. **d**, Experimental design for anterograde labeling of SuM-recipient MS neurons by EGFP, and labeling of vGlut2^+^ neurons by injection of DIO-mCherry in vGlut2-Cre mice. **e**, Representative confocal images showing mCherry (vGlut2+) and EGFP anterograde) double-labeled cell bodies in MS. **f**, Summary of co-labeling proportion of SuM-recipient and vGlut2+ neurons in MS, *n* = 3 mice. **g**, Experimental design for anterograde labeling of SuM-recipient MS neurons by EGFP, and immunohistochemistry (IHC) for CHAT in MS. **h**, Representative confocal images showing CHAT (IHC) and EGFP (anterograde) double-labeled cell bodies in MS. **i**, Summary of co-labeling proportion of SuM-recipient and CHAT^+^ neurons in MS, n = 3 mice.

**Figure5–figure supplement 2.**
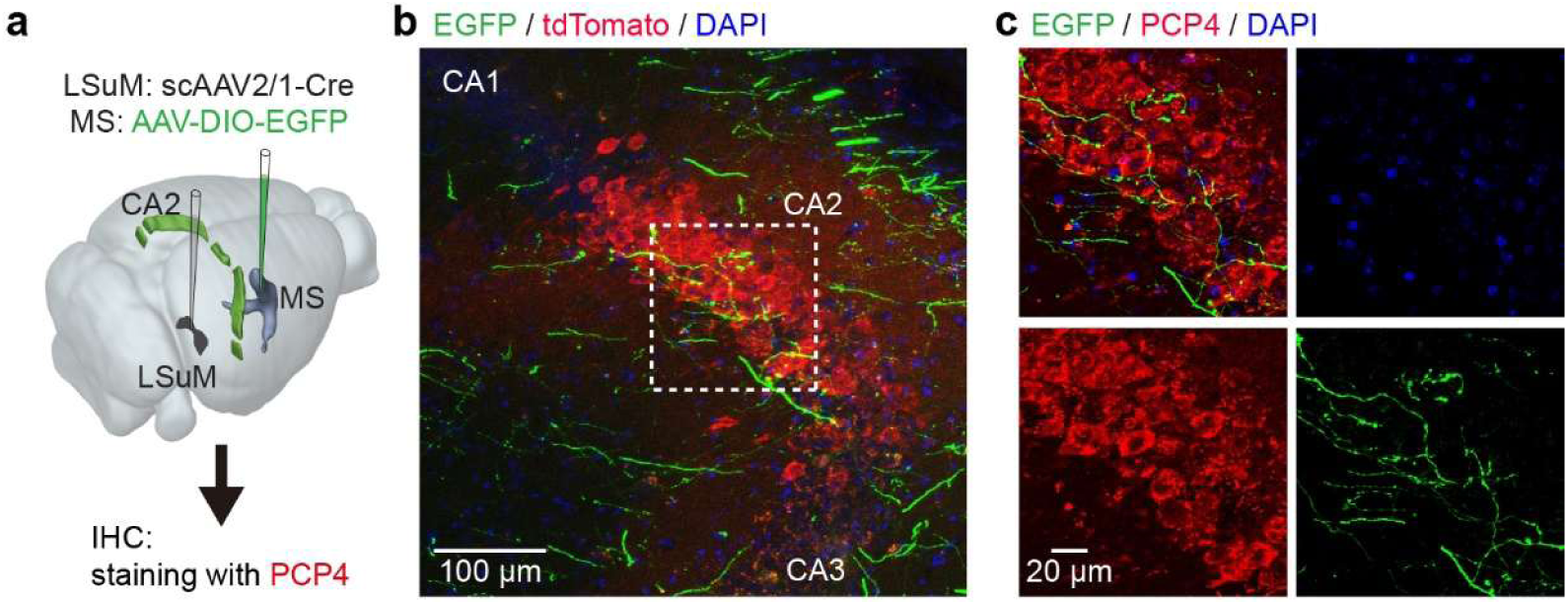
Anatomical connectivity from SuM-recipient MS neurons to CA2. **a**, Experimental design for anterograde labeling of EGFP in SuM-recipient MS neurons. **b**, Representative confocal images showing EGFP-labeled axonal terminals in the CA2 region. **c**, Magnified view of the boxed area in **b**.

## Methods

### Animals

Adult male C57BL/6CJ mice (3–5 months old) were used for tracing, recording and behavioral experiments. vGlut2-Cre (Stock No. 016963), vGAT-Cre (Stock No. 028862) and LSL-H2B-GFP (Stock No. 036761) transgenic mice were used for morphology experiments. Mice were group-housed (4-5 per cage) unless implanted with optical fibers or tetrodes, in which case they were singly housed. All animals were maintained on a 12 h light/dark cycle (lights on at 7:00) with ad libitum access to food and water. All experimental procedures were approved by the Third Military Medical University Animal Care and Use Committee and conducted in accordance with their guidelines and protocols.

### AAVs

The following adeno-associated virus (AAV) constructs were used: AAV2/8-EF1α-EGFP (titer: 1.49 × 10¹³ vp/mL), AAV2/2Retro-Syn-GCaMP6m (titer: 1.92 × 10¹³ vp/mL), AAV2/2Retro-Syn-H2B-mRuby3 (titer: 1.92 × 10¹³ vp/mL), AAV2/2Retro Plus-Syn-Cre (titer: 1.92 × 10¹³ vp/mL), scAAV2/1-Syn-Cre (titer: 1.92 × 10¹³ vp/mL), scAAV2/1-Syn-Flpo (titer: 1.92 × 10¹³ vp/mL), AAV2/9-EF1α-DIO-hChR2-mCherry (titer: 3.67 × 10¹³ vp/mL), AAV2/9-Syn-DIO-hGtACR1-mCherry (titer: 1.43 × 10¹³ vp/mL), AAV2/9-Syn-DIO-mCherry (titer: 1.23 × 10¹³ vp/mL), AAV2/9-Syn-DIO-EGFP (titer: 1.23 × 10¹³ vp/mL), AAV2/9-Syn-fDIO-EGFP (titer: 1.23 × 10¹³ vp/mL). All AAVs were purchased from Taitool Bioscience Co., Ltd. (Shanghai, China).

### Surgical Procedures

For all surgeries, mice (2-3 months old) were anesthetized with isoflurane (3% induction, 1-2% maintenance in oxygen) and secured in a stereotaxic frame. Body temperature was maintained at 37.5-38°C using a heating pad. Following surgery, mice were returned to warmed cages for full recovery and received daily intraperitoneal injections of dexamethasone sodium phosphate (1 mg/mL; 0.1 mL per 10 g body weight) and ceftriaxone sodium (50 mg/mL; 0.1 mL per 10 g body weight) for three consecutive days to mitigate post-operative inflammation.

For stereotactic injection, viral vectors were pressure-injected using a glass micropipette (tip diameter: 10-20 µm) into target brain regions at the following coordinates (relative to bregma, mm): SuM, AP: -2.8, ML: ±0.5, DV: -5.0 (from dura); MS, AP: 1.0, ML: 0.5, 5° angle toward the midline, DV: 3.8; CA2, AP: -1.8, ML: ±2.45, DV: -1.55; DG: AP: -1.8, ML: ±1.1, DV: -1.85. Injections were unilateral for tracing/recording experiments and bilateral for behavioral experiments. Viruses were allowed a minimum expression period of 4 weeks prior to subsequent experiments.

For optical fiber implantation, mice expressing GCaMP6m in SuM-MS projection were used. A 200-µm diameter, 0.53 NA optical fiber (Doric Lenses, MFP_200/230/900-0.53) was glued into a metal cannula (ID 0.51 mm, OD 0.82 mm) with a flat-polished end. The prepared fiber probe was lowered through a cranial window above the lateral SuM (AP: -2.8 mm, ML: 0.5 mm) to the depth of 5.0 mm. The cannula was secured to the skull using blue light-curing dental cement (Tetric EvoFlow). Additional dental cement and cyanoacrylate glue provided reinforcement. The outer surface was coated with black acrylic paint to prevent light leakage.

For EEG and EMG electrode implantation, three EEG screws were placed over the skull: two above the frontal cortex (AP: +1.5 mm, ML: ±1.5 mm) and one above the parietal cortex (AP: -3.0 mm, ML: +3.0 mm). Two EMG wires were inserted into the nuchal muscles. All electrodes were secured using blue light-curing dental cement, followed by reinforcement with additional dental cement and cyanoacrylate glue.

For optrode implantation, mice expressing ChR2 in SuM-MS projection were used. A custom optrode (Yang et al., 2023) was mounted on a microdrive, consisting of four linearly arranged 25-μm tungsten wire tetrodes spaced at 200-μm intervals, with tips extending approximately 500 μm beyond a 200-μm diameter, 0.37 NA optical fiber. The optrode was initially positioned above the lateral SuM (AP: −2.8 mm, ML: 0.5 mm, DV: −4.7 mm). EEG/EMG electrodes were implanted as described previously. Following one week of recovery, the optrode was gradually advanced to the target depth (−5.0 mm). Finally, an electrolytic lesion (100 μA, 10 s) was applied to mark the recording site for subsequent histological verification.

For fiber ferrule implantation, mice expressing GtACR1-mCherry or mCherry bilaterally in SuM-MS or MS-CA2 projection were used. Fiber ferrules (200 µm diameter, 0.37 NA) were implanted above SuM (AP: -2.8 mm, ML: ±2.0 mm, DV: -4.6 mm, angled 20° towards midline) or MS (AP: 1.0 mm, ML: 0.5 mm, 5° angle toward the midline, DV: 3.8 mm).

### Fiber Photometry Recording

Mice implanted with optical fibers above lateral SuM with SuM-MS projecting neurons expressing GCaMP6m were used. Ca²⁺ signals, EEG/EMG signals, and behavioral videos were recorded simultaneously across sleep-wake cycles. Signals were acquired at 2000 Hz for Ca²⁺, 200 Hz for EEG/EMG, and 25 Hz for video. All data streams were synchronized offline using event markers.

### Behavioral Tasks

Mice were handled for 5 min/day for continuous 3 days prior to behavioral habituation. All tasks were conducted between 7:00 - 8:00 am. Optogenetic silencing was performed using a 473 nm laser (controlled via LabVIEW) delivered bilaterally through implanted optical fibers, with a light intensity of ∼3 mW at each ferrule tip. For REM sleep-specific silencing, light delivery was triggered immediately upon the onset of a REM episode (identified in real-time via EEG/EMG) and remained continuously on until the episode ended. This paradigm was applied to both GtACR1-expressing and mCherry control groups during the 8-hour post-training consolidation periods of the social and contextual fear memory tasks. In an additional GtACR1 control group, light was delivered during NREM sleep or quiet wakefulness, with the cumulative light duration matched to that of the REM-silenced group.

### Three-chamber Social Test

The three-chamber social test was employed to assess social memory, as previously described (Qin et al., 2022). The apparatus consisted of a rectangular box (75 cm × 50 cm × 40 cm) divided into three equally sized chambers (each 25 cm × 50 cm) by transparent acrylic partitions. Two transparent cylinders (diameter: 12 cm) with wall air holes were used to contain stimulus mice. On day 1, the subject mouse was habituated to the apparatus during a 10-min session with both cylinders empty. On day 2 (training), the mouse explored for 20 min with one cylinder containing a novel stimulus mouse and the other remaining empty. EEG/EMG recording began immediately after training and continued for the subsequent 8 h. NREM and REM sleep episodes were manually identified based on the scoring criteria detailed later for silencing interventions. In the GtACR1 silencing group and the mCherry control group, blue light was delivered continuously to the SuM starting ∼6 s after REM sleep detection and persisting until the end of each REM episode. In the GtACR1 control group, blue light was administered during NREM sleep or quiet wakefulness for a cumulative duration matched to the total REM-light time in the experimental group. On day 3 (test), the subject mouse was reintroduced to the apparatus for a 20-min session, during which one cylinder contained the now-familiar mouse and the other contained a novel mouse. Time spent investigating each cylinder was recorded and the social preference index was calculated as:

Social preference index = (Time_Novel_ - Time_Familiar_) / (Time_Novel_ + Time_Familiar_).

### Contextual Fear Conditioning

The task was conducted to test contextual memory as previously described (Boyce et al., 2016). On the first day (training), after a 260-s baseline period, each subject mouse received three pairings of a conditioned auditory stimulus (CS: 8964 Hz pure tone, 20 s, 70 dB SPL) followed by an unconditioned stimulus (US: 2 s, 0.6 mA footshock), with 180 s intervals between pairings. EEG/EMG were recorded for 8 h immediately after training, and optogenetic silencing was applied as described earlier. Fear memory was tested 24 h after training. Contextual memory was assessed by re-exposing the mouse to the training context for 10 min. Freezing level was quantified based on the absence of body movement.

### Sleep Analysis

EEG and EMG signals were first filtered (0.5–30 Hz and 10–70 Hz, respectively). Sleep-wake states were then determined based on visual inspection of the following features: wakefulness exhibited low-amplitude, high-frequency EEG with elevated EMG; NREM sleep showed high-amplitude delta (0.5–4 Hz) EEG and low EMG tone; REM sleep was marked by theta-dominated EEG (4–10 Hz) with absent tonic EMG activity.

### Histology

Mice were transcardially perfused with 4% paraformaldehyde (PFA) in PBS. Brains were post-fixed (4% PFA, 24h), cryoprotected (15% sucrose, 24h), and sectioned coronally (50 µm). Sections were stained with DAPI and imaged (confocal: Zeiss LSM 700; widefield: Olympus BX51). For immunohistochemistry, primary antibodies against PCP4 (rabbit polyclonal, 1:500; Sigma-Aldrich, Cat# HPA005792) and ChAT (goat polyclonal, 1:500; Millipore, Cat# AB144P) were applied.

### Quantification and Statistical Analysis

For fiber recording data, Signals were low-pass filtered using a Savitzky–Golay filter (50-point window, 3rd-order polynomial). ΔF/F was computed as (F - F_baseline_) / F_baseline_, where F_baseline_ represents the mean fluorescence over the period that without Ca^2+^ transients. The area under the curve (AUC) of ΔF/F traces was quantified for each defined sleep-wake state.

For single-unit data, extracellular electrophysiological signals were preprocessed (Qin et al., 2018) and spike-sorted with the MClust toolbox(Schmitzer-Torbert and Redish, 2004) based on waveform features. Firing rates were calculated using 2-s sliding windows (100-ms step). Units were classified according to their firing patterns.

For spectral analysis, CA1 LFP and EEG power spectra were obtained via fast Fourier transformation by custom MATLAB code (frequency resolution 0.15 Hz, (Qin et al., 2022). Mean power density in the delta (1–4 Hz), theta (6–12 Hz), alpha (12–15 Hz), and beta (15–30 Hz) bands (Billwiller et al., 2020, Boyce et al., 2016) was compared across 4-s windows before, during, and after optogenetic silencing.

For statistics, all analyses were performed in MATLAB and SPSS 22.0. Normality and homogeneity of variance were tested before selecting appropriate tests. If assumptions were met, parametric tests were applied (paired/unpaired t-tests, 1-way ANOVA with Bonferroni post-hoc, or RMs 2-way ANOVA with Bonferroni post-hoc). Otherwise, non-parametric alternatives were used (Wilcoxon signed-rank test, Wilcoxon rank-sum test, or Friedman test with Bonferroni post-hoc comparison). Data are reported as mean ± SEM.

## Acknowledgements

The authors are grateful to Ms. Jia Lou for help in composing and layout editing of the figures. This work was supported by grants from the National Natural Science Foundation of China to X.C. (32430044, 32127801) and H.Q. (32200838), the National Key R&D Program of China to X.C. (2021YFA0805000), the Chongqing Natural Science Foundation Project to H.Q. (CSTB2024NSCQ-QCXMX0043), and the Jiangsu Provincial Big Science Facility Initiative to H.J. (BM2022010).

## Ethics

All animal experiment procedures were approved by the Animal Care and Use Committee of the Third Military Medical University (Approval number: AMUWEC20211064). Experiments were conducted strictly in adherence to the National Institutes of Health Guidelines for Care and Use of Laboratory Animals.

## Author contributions

Conceptualization, H.Q., X.C. and J.H; Investigation, T.J., W.J., and M.L.; Formal analysis, T.J., X.L., K.Z., S.L., and C.Z.; Visualization, T.J., C.H., H.J., Y.W. and H.Q.; Writing–original draft, H.Q.; Writing–review and editing, H.Q., J.H. and X.C.; Supervision, H.Q., J.H. and X.C.

## Declaration of interests

The authors declare no competing interests.

## Notes

### Competing Interest Statement

The authors have declared no competing interest.

### Summary of Updates

Added viral tracing data on neurotransmitter phenotypes (Figure 1-figure supplement 1; Figure 5-figure supplement 1). Added anterograde tracing confirming SuM-recipient MS neuron-CA2 connectivity (Figure 5-figure supplement 2). Added histology validating injection and fiber sites for CA2, SuM and MS (Figures 4, 5). Added a Discussion limitation on lack of causal proof for the full SuM-MS-CA2 chain. Replaced LSD with Bonferroni tests and added test statistics. Revised figure titles/legends to specify recording modalities. Corrected figure citations and added Figure 3c, 3d citations. Detailed optogenetic silencing light parameters. Expanded Discussion on mechanisms, regional heterogeneity and future directions.

